# Genomic Profiling of Estrogen-Related Receptor Identifies Adult Adipocyte-Specific Targets in *Drosophila*

**DOI:** 10.64898/2026.07.29.741485

**Authors:** Jun Yin, Douglas B. Rusch, Jitika S. Bhatta, David M. Merritt, Lesley N. Weaver

## Abstract

Adipose tissue is a key metabolic organ for carbohydrate and lipid metabolism. Multiple nutrient sensing pathways and metabolic enzymes operate in adipocytes to control energy production and lipid mobilization in response to physiological conditions. Important regulators of glycolytic and lipid metabolism in many species are members of the estrogen-related receptor (ERR) family of nuclear receptors. We previously showed that ERR is required for transcriptional regulation of glycolytic and pentose phosphate pathway enzymes in adult *Drosophila* females, consistent with what is observed in *ERR* mutant males and larvae. However, the cell-type specific targets of ERR in adipose and other tissues have not been fully elucidated. Here, using a modified targeted DamID approach (NanoDam), we identified ERR occupancy specifically in adult adipocytes at the loci that encode genes involved in glycolysis, the pentose phosphate pathway, and fatty acid metabolism. Overall, our results predict that ERR centrally functions in adipose tissue to regulate multiple metabolic pathways.

**SUMMARY:** Identification of genomic loci bound by proteins can be difficult to isolate in tissues composed of multiple cell types or if an antibody is unavailable. Here, we used a genomic approach called NanoDam to identify adult adipocyte-specific targets of the nuclear receptor, estrogen-related receptor (ERR), in *Drosophila* males and females. We find that ERR targets in both sexes are enriched for genes encoding enzymes involved in glycolysis and the pentose phosphate pathway, suggesting that ERR targets are similar in adult adipocytes as in larval fat. Our results will be of interest for scientists studying nuclear receptors, metabolism, and sexual dimorphism.

## INTRODUCTION

Multicellular organisms are composed of complex tissues, and organ function is dependent on the coordinated effort between tissues to maintain energy homeostasis (Castillo-Armengol et al. 2019; Priest and Tontonoz 2019). Adipose tissue is a major nutrient sensing endocrine organ, functioning as a key site for energy storage and lipid mobilization (Corvera et al. 2026; Sakers et al. 2022). Adipocytes synthesize triglycerides from glucose for storage in lipid droplets, as well as breakdown and distribute fatty acids to the liver in response to fasting conditions (An et al. 2023; Konige et al. 2014; Luo and Liu 2016). The *Drosophila* fat body (composed of adipocytes and hepatocyte-like oenocytes) functions as a compound tissue akin to mammalian adipose and liver tissues (Huang and Perrimon 2026; Li et al. 2019). Thus, it serves as an excellent model for understanding inter-organ signaling and metabolism (Chatterjee and Perrimon 2021; Musselman and Kuhnlein 2018; Trinh and Boulianne 2013). Disruptions in adipocyte homeostasis are linked to pathologies such as obesity, diabetes, and inflammation (Chait and den Hartigh 2020; Lin and Jensen 2025; Wang et al. 2025). Therefore, determining the key regulators that function in adipose tissue and understanding their direct metabolic targets may provide insight for future therapeutic interventions.

Nuclear receptors are ligand-dependent transcription factors that control many biological processes, including metabolism (Dixon et al. 2021; He et al. 2025; Jacobi et al. 2012; Li et al. 2025; Scholtes and Giguere 2022). For example, peroxisome proliferator-activated receptor (PPAR) β/δ null mutant mice have increased adipocyte lipolysis (Kang et al. 2008), suggesting a role in fatty acid oxidation. In addition, in response to high sugar conditions, liver X receptor (LXR) increases expression of glycolytic lipogenic enzymes to regulate glucose levels (Bindesboll et al. 2015). Nuclear receptors are also the targets of endocrine disruptors (La Merrill et al. 2025; La Merrill et al. 2020; Lu et al. 2026; Tabb and Blumberg 2006; Toporova and Balaguer 2020), therefore determining their tissue-specific targets may provide insight to the mechanisms disrupted in metabolic syndrome disorders, as well as the biological pathways perturbed by xenobiotics.

Estrogen-related receptors (ERRs) are orphan nuclear receptors that control metabolism and energy homeostasis in multiple species (Audet-Walsh and Giguere 2015; Giguere 2008; Spinelli et al. 2024; Tripathi et al. 2020). For example, ERR α and γ are required in mammalian adipocytes to regulate metabolic processes such as glycogen metabolism (Fu et al. 2025), fatty acid oxidation (Dixen et al. 2013), mitochondrial biogenesis (Schreiber et al. 2004; Singh et al. 2018), and thermogenesis (Ahmadian et al. 2018; Brown et al. 2018; Dixen et al. 2013; Villena et al. 2007). Furthermore, ERRα null mice have reduced body fat and are resistant to high fat diet induced obesity (Luo et al. 2003). In *Aedes aegypti* (the yellow fever mosquito), *ERR* expression is induced in an ecdysone-dependent manner after a blood meal to promote carbohydrate and lipid metabolism (Geng et al. 2024). Furthermore, *ERR* is highly expressed in *Bombyx mori* (the domestic silkworm) fat bodies and binds to the promoters of genes involved in lipid, amino acid, and carbohydrate metabolism (Hou et al. 2025).

In *Drosophila*, loss of the sole *ERR* homolog decreases glycolytic enzyme transcription (Beebe et al. 2020; Tennessen et al. 2011; Zike et al. 2025) and diminishes lipid accumulation in adipocytes (Fasteen et al. 2025). We recently showed that whole body *ERR* knockout in adult females differentially regulates transcripts of enzymes involved in glycolysis and the pentose phosphate pathway, with some differentially expressed genes showing sexual dimorphism (Fleck et al. 2026); however, the tissue-specific ERR targets mediating these changes are unknown. The conserved defects in carbohydrate and lipid metabolism in the absence of *ERR* in multiple species suggest that ERR is a central metabolic regulator. Despite these advances, it remains unclear whether the observed changes in mRNA expression levels in *ERR* mutants is due to direct ERR regulation of those genes.

Identification of cell-type specific transcription factor targets can be challenging in tissues that contain different cell identities. Targeted DNA adenine methyltransferase identification (TaDa) approaches allow for cell-type specific identification of transcription binding in multiple organisms (Cheetham et al. 2018; Katsanos and Barkoulas 2022; Southall et al. 2013) by fusing the Dam methytransferase from *E. coli* (van Steensel and Henikoff 2000) to the transcription factor of interest. This technique is beneficial because targets are identified without requiring organ dissection or cell isolation, and does not require large sample sizes or specific antibodies. NanoDam is a modified targeted TaDa approach that enables cell-specific identification of binding sites of endogenously GFP-tagged transcription factors using a Dam methyltransferase construct fused to a GFP nanobody that is expressed only in specific cell types (Tang et al. 2022a; Tang et al. 2022b). In *Drosophila*, the NanoDam transgene is under control of an upstream activator sequence (UAS) that is activated by the Gal4 driver that is under control of a cell-type specific promoter. When crossed with flies containing the GFP-tagged transcription factor, site-specific methylation of adenines in GATC sequences only occurs where expression of the NanoDam and transcription factor overlap. This approach is highly advantageous because large tissues quantities do not need to be dissected (or cell types sorted), an antibody is not required, and cell-specific targets are identified through enrichment of methylated sites.

In this study, we investigated which genes are directly regulated by ERR in adult *Drosophila* male and female adipocytes. We used NanoDam (Tang et al. 2022a; Tang et al. 2022b), a modified targeted DamID approach that recruits the Dam methylase to endogenously tagged proteins using nanobodies, to identify targets of an endogenously GFP-tagged ERR protein specifically in adult male and female adipocytes. We found that ERR binds to the loci of a core set of enzymes involved in glycolysis, the pentose phosphate pathway (PPP), and β-oxidation in the adipocytes of both males and females. Whereas ERR binding in male adipocytes was enriched for genes involved in fatty acid and carbohydrate metabolism, ERR occupancy in females showed enrichment for glycogen and carbohydrate metabolic genes. Our results herein predict that ERR activity is required specifically in the adipocytes of both sexes to regulate central metabolic pathways.

## EXPERIMENTAL PROCEDURES

### Drosophila strains and culture

*Drosophila* strains were maintained on BDSC cornmeal food (15.9 g/L inactive yeast, 9.2 g/L soy flour, 67.1 g/L yellow cornmeal, 5.3 g/L agar, 70.6 g/L light corn syrup, 0.059 M propionic acid) at 22-25°C. Due to proper oogenesis requiring a rich protein source, all females used in experiments were maintained on BDSC cornmeal food supplemented with wet active yeast paste and flipped to fresh food daily. Experimental males were maintained on BDSC cornmeal food and flipped to fresh food each day. Although males are routinely kept on fly food without wet yeast supplementation, it is possible observed sex differences are due to changes in diet. Flies were kept at a 12 hr/12 hr light/dark cycle at all temperatures unless otherwise noted. The previously described *UAS-NanoDam* (Tang et al. 2022a) transgene was used to methylate adenine residues in GATC sequences. The previously described *Gal4* and *Gal80* transgenes were used, including *3.1Lsp2-Gal4* [RRID:BDSC_98128, (Armstrong et al. 2014; Lazareva et al. 2007)] and *tubGal80^ts^* [RRID:BDSC_7019, (McGuire et al. 2003)] and were obtained from the Bloomington *Drosophila* Stock Center (BDSC, bdsc.indiana.edu). Additional stocks obtained from the BDSC include *y^1^ w^1^* (RRID:BDSC_1495) and *w^1118^; PBac{ERR-GFP.FSTF}VK00037* (RRID:BDSC_38638, (Venken et al. 2009)). Lines carrying multiple genetic elements were generated by standard crosses. Balancer chromosomes and genetic elements are described on Flybase (flybase.org), which was used as a reference tool throughout this study (Ozturk-Colak et al. 2024).

### Adult Adipocyte-Specific ERR NanoDam

Females and males of genotypes *y w; tubGal80^ts^/+; 3.1Lsp2-Gal4/UAS-NanoDam tubGal80^ts^* and *y w; tubGal80^ts^/ERR::eGFSTF; 3.1Lsp2-Gal4/UAS-NanoDam tubGal80^ts^* (for adult adipocyte-specific ERR-NanoDam) were raised at 18°C [the permissive temperature for Gal80^ts^ (McGuire et al. 2003)] to prevent NanoDam DNA methylation during development. Zero-to-two-day-old females were maintained at 18°C for one to two days with *y^1^ w^1^* males and then switched to 29°C for 14 hours. Males of each genotype were housed separately under the same temperature conditions and flipped to fresh food without wet yeast paste each day prior to NanoDam induction. 10 whole males or females of each genotype were flash frozen in liquid nitrogen and stored at -80°C. Four biological replicates were generated for each experiment. Genomic DNA was extracted using the Purgene Tissue Kit (Qiagen) according to manufacturer’s instructions. Methylated DNA enrichment and library preparation were carried out as previous described (Tang et al. 2022a; Tang et al. 2022b), with slight modifications. Libraries were sequenced on an Illumina Nextseq2000 platformed with paired-end 2x61 bp sequencing reads. Greater than 15 million reads were obtained for each library. Samples were trimmed for quality and adapters using fastp v0.21.0 (Chen et al. 2018) using default parameters. The trimmed reads were then mapped with bowtie2 v2.5.4 (Langmead and Salzberg 2012) (non-default parameters: --local) to the *Drosophila* genome (r6.42) in local mode to exclude special adapters that were not efficiently removed during library preparation. Counts were determined by identifying properly paired inserts that spanned a region less than 1,000 bps; any DpnI restriction fragment that the insert overlapped was incremented by one. The resulting counts table was analyzed using the R language package DESeq2 (Love et al. 2014; Southall et al. 2013) to identify differentially abundant DpnI fragments between the experimental and control samples. A DpnI fragment overlapped with a gene if it was within 1 kb of the start and end site of the gene. Differentially abundant fragments with an adjusted *P* value less than 0.001 and log2 fold change ≥ 2 (indicating at least a 4-fold increase in methylation compared to the NanoDam only control) were considered significant. Significant negative log2 fold change values (indicating decreased binding) were observed in ∼10% of the samples as previously described (Southall et al. 2013), and were considered stochastic noise. Mapped reads and inserts were visualized in JBrowse (Buels et al. 2016). Gene ontology (GO), KEGG pathway, and REACTOME pathway enrichment of genes surrounding significant binding loci were analyzed using PAthway, Network and Gene-set Enrichment Analysis (PANGEA) (Hu et al. 2023).

## RESULTS AND DISCUSSION

### Genomic profiling using NanoDam identifies over 900 adipocyte-specific ERR binding sites in males and females

ERR is a central regulator of carbohydrate metabolism in *Drosophila* larvae and adult male fat bodies (Beebe et al. 2020; Fasteen et al. 2025); however, the direct adipocyte-specific ERR targets during development and between the sexes are unknown. The *Drosophila* fat body consists of both adipocytes and closely associated hepatocyte-like oenocyte cells (Li et al. 2019), making identification of cell-type specific ERR targets challenging. We therefore used NanoDam to identify the direct targets of ERR in adult female and male adipocytes (**Figure 1a**). We expressed NanoDam for 14 hours in adult male and female adipocytes using the previously described adipocyte-specific *3.1Lsp2-Gal4* driver (Armstrong et al. 2014; Lazareva et al. 2007) with *tubGal80^ts^* (McGuire et al. 2003) (hereafter referred to as *3.1Lsp2^ts^*) in the presence of an endogenously C-terminal eGFP-tagged ERR bacterial artificial chromosome (*ERR::eGFSTF*). We previously showed that the *ERR::GFSTF* BAC rescues germline stem cell loss in adult *Drosophila* females (Zike et al. 2025), suggesting that the multiple tags (eGFP, FlAsh, StrepII, Tev, and FLAG) do not interfere with ERR activity. Compared to the NanoDam alone negative control, ERR-NanoDam in adult female adipocytes identified over 950 regions in the genome occupied by ERR representing more than 1,000 unique genes (**Figure 1b; Supplementary Table 1**). In contrast, ERR occupancy was observed at 1,580 regions in male adipocytes, representing 1480 unique genes (**Figure 1c, Supplementary Table 2**). In both datasets, we noted that some identified regions spread across multiple genes, resulting in more genes that are potentially bound by ERR than genomic regions. Furthermore, we also observed that some regions indicated decreased ERR enrichment compared to the NanoDam-alone as previously observed for TaDa experiments (Southall et al. 2013), suggesting these regions contained more methylation in the NanoDam alone negative control. We therefore attributed decreased ERR-NanoDam enrichment as stochastic noise, as negative binding only accounted for ∼10% of significant changes of binding in both datasets.

**Figure 1.**
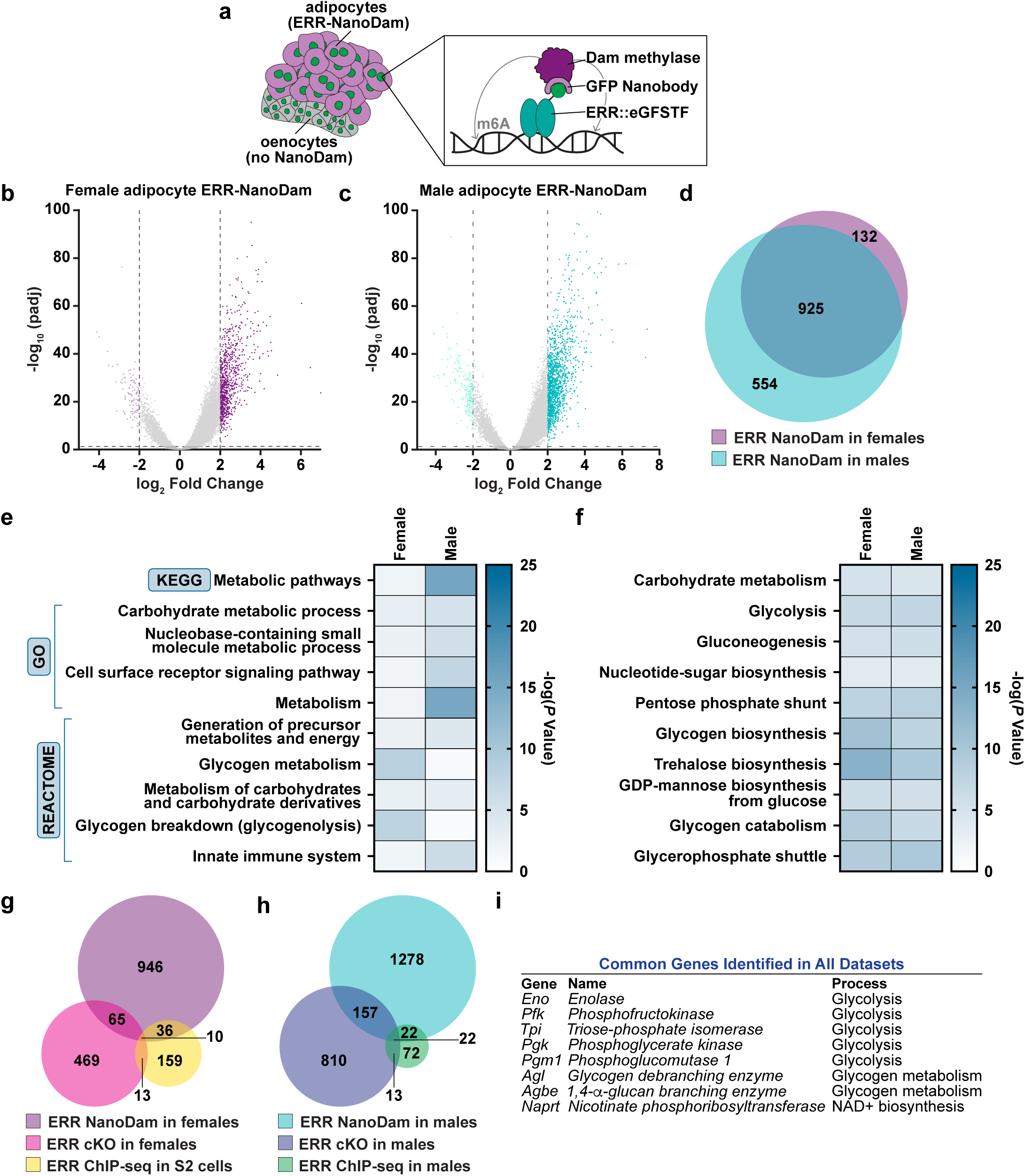
Adipocyte-specific ERR NanoDam identifies genomic regions bound in adult females and males. **(a)** Cartoon schematic of NanoDam (Tang et al. 2022a). **(b, c)** Volcano plots of adipocyte ERR-NanoDam in females (b) and males (c) indicating regions of the genome positively bound by ERR based on a log2 fold change ≥ 2 (darker colors). Negative log2 Fold Change values ≤ -2 (lighter colors) were excluded as stochastic noise. **(d)** Venn diagram of genes located near ERR bound regions of the *Drosophila* genome in adult female and male adipocytes. **(e)** Heatmap showing the top 10 gene sets based on the adjusted *P*-value from PANGEA KEGG, GO, and REACTOME enrichment. **(f)** Heatmap showing the top 10 gene sets based on the adjusted *P*-value using “Fly Metabolic Pathway” analysis. **(g)** Venn diagram of genes identified in female-specific adipocyte ERR-NanoDam, *ERR* adult female mutant whole-body RNA-seq, and S2 cell ERR ChIP-seq datasets. **(h)** Venn diagram of genes identified in male-specific adipocyte ERR-NanoDam, *ERR* adult male mutant whole-body RNA-seq, and whole-body male ERR ChIP-seq datasets. **(i)** Table showing the common genes identified from all datasets in (g) and (h).

Comparison of the genes in which ERR was bound showed that 925 genes were common between females and males (**Figure 1d**). Pathway, Network, and Gene-set Enrichment Analysis (PANGEA) (Hu et al. 2023) using the KEGG, REATOME and gene ontology selections showed that the common genes near ERR binding were enriched for metabolic pathways, carbohydrate metabolic processes, and glycogen metabolism (**Figure 1e, Supplementary Table 3**). Furthermore, selection of the “Fly Metabolic Pathway” setting in PANGEA identified enrichment for carbohydrate metabolism, glycolysis, trehalose biosynthesis, and the pentose phosphate shunt (**Figure 1f, Supplementary Table 4**). Analysis of the 132 female-specific predicted targets using the “Fly metabolic pathway” function in PANGEA showed significant enrichment for protein glycosylation and dolichol-linked oligosaccharide biosynthesis (**Supplementary Table 5**); whereas, the male-specific predicted targets were enriched for metabolic pathways, including, carbohydrate metabolism, gluconeogenesis, and the malate-asparate shuttle (**Supplementary Table 6**).

We also compared the female adipocyte ERR-NanoDam dataset to our previous RNA-seq dataset that identified differentially expressed genes in whole-body *ERR* mutant adult females (Fleck et al. 2026) and an S2 cell ERR ChIP-seq dataset (as a female-specific dataset is unavailable; (Kovalenko et al. 2019)). Ten genes were identified in all three datasets (**Figure 1g, Supplementary Table 7**), indicating these genes are likely direct ERR targets regardless of cell type and developmental stage. Furthermore, analysis of male adipocyte ERR-NanoDam with published adult *ERR* mutant whole-body RNA-seq and adult male ERR ChIP-seq datasets (Beebe et al. 2020) identified 22 common genes (**Figure 1h, Supplementary Table 8**). Comparison of all six referenced datasets identified 8 genes (**Figure 1i**) with known roles in metabolic pathways, indicating these genes are likely direct ERR targets regardless of cell type, developmental stage, and sex. These results are consistent with the roles of ERR in regulating carbohydrate metabolism, glycolysis, and the PPP in insects and mammals (Beebe et al. 2020; Dixen et al. 2013; Fasteen et al. 2025; Fleck et al. 2026; Fu et al. 2025; Geng et al. 2024; Hou et al. 2025; Luo et al. 2003; Tennessen et al. 2011; Zike et al. 2025). Collectively, our results suggest that ERR adipocyte metabolic control is conserved between species, at different developmental stages, and between the sexes.

### ERR binds to genomic loci that encode glycolytic enzyme genes in adult Drosophila female and male adipocytes

Our metabolic pathway analysis indicated genes involved in the breakdown of glucose to pyruvate (glycolysis, **Figure 2a**) are bound by ERR in both female and male adipocytes. ERRs are known regulators of glycolysis, and knockdown of *ERR* in larval adipocytes significantly decreases transcript expression of glycolytic enzymes (Fasteen et al. 2025). To determine which glycolytic genes may be direct targets of ERR in adipocytes, we analyzed binding of each enzyme in males and females (**Figure 2b and c**). ERR showed genomic binding in both male and female adipocytes for seven of 13 glycolytic enzymes; whereas males had higher binding for an additional 3 genes (*Pgi*, *Gapdh1*, and *Gapdh2*) (**Figure 2c**). Neither male or female adult adipocytes showed significant occupancy for *Hex-C*, *Ald*, or *Pglym78*. These results suggest that ERR likely binds to the loci of genes encoding glycolytic enzymes in adult *Drosophila* adipocytes.

**Figure 2.**
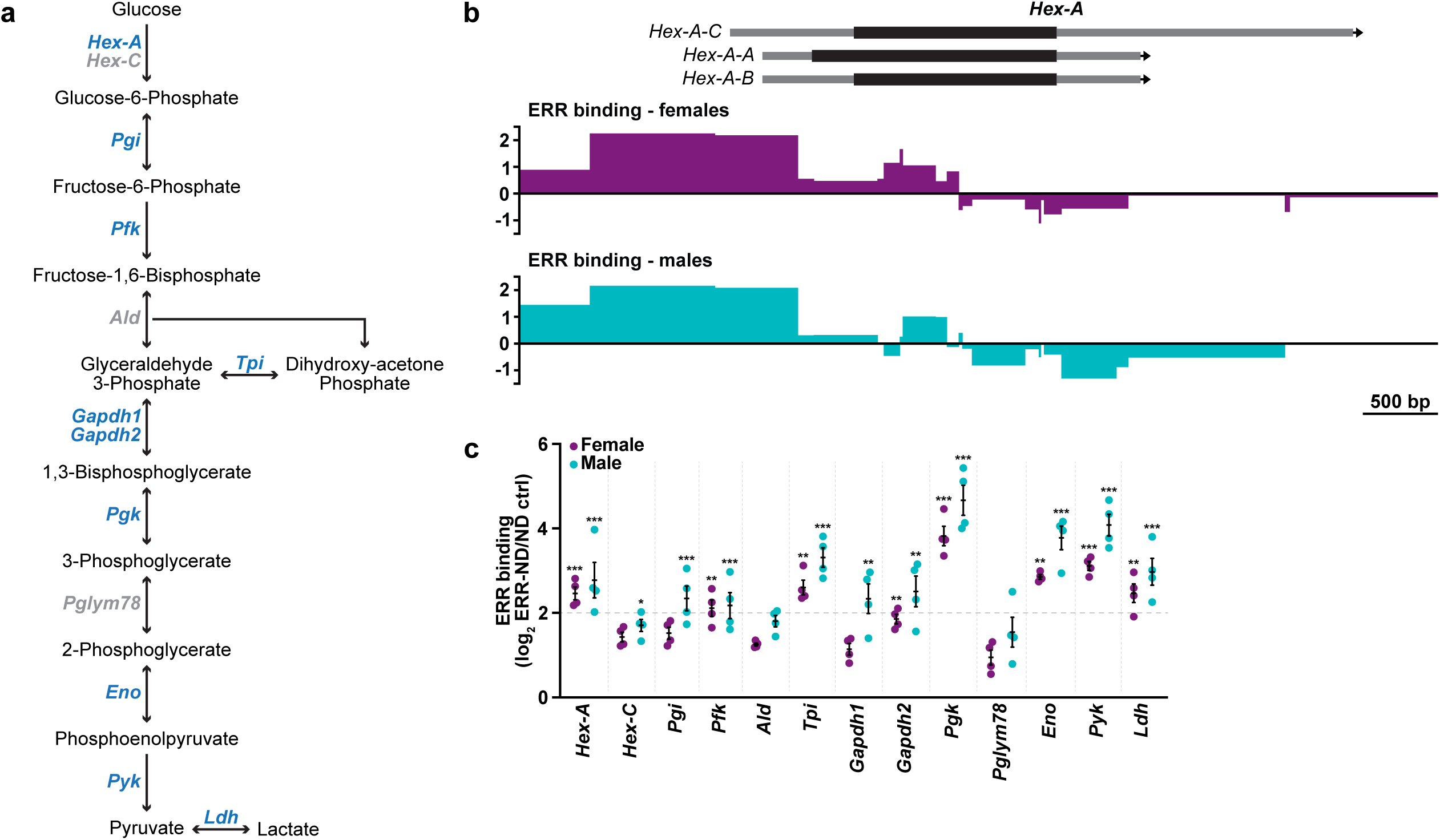
Glycolytic enzyme loci are occupied by ERR in adult adipocytes. **(a)** Diagram of glycolysis showing the intermediates and key enzymes. Enzymes bound by ERR in adult female or male adipocytes are bolded in blue font. **(b)** Representation of the *Hex-A* genomic locus and ERR binding in female (purple) and male (teal) adipocytes indicating positive binding of ERR based on a log2 fold change ≥2. Negative log2 fold change values ≤ -2 were considered stochastic noise. **(c)** Quantification of ERR binding (ERR-ND) in adult female and male adipocytes at glycolytic enzyme loci compared to the NanoDam alone negative control (ND ctrl). Data from four independent experiments shown as mean ± SEM. Dotted line indicates threshold ratio of ERR-ND/ND ctrl = 2. Values ≥ 2 were considered significant binding as a potential ERR-NanoDam target. \**P* < 1x10^-10^, \*\**P* < 1x10^-15^, \*\*\**P* < 1x10^-20^; Wald test.

Analysis comparing adipocyte ERR-NanoDam with previously published RNA sequencing (RNA-seq) datasets from *Drosophila* whole body *ERR* conditional knockouts (Beebe et al. 2020; Fleck et al. 2026) show overlap with five genes (*Pfk*, *Tpi*, *Pgk*, *Eno*, and *Pyk*) in females; whereas, all putative ERR-NanoDam candidates except *Hex-A* and *Pyk* were significantly downregulated in males, suggesting ERR may regulate differentially expressed glycolytic genes in adult adipocytes. Furthermore, ERR-bound genes in adult adipocytes strongly align with genes significantly downregulated in larval adipocytes from *ERR* mutants (10 of 12 overlapping genes) (Fasteen et al. 2025). While mammalian ERRs are preferentially expressed in brown adipose tissue to regulate genes involved in mitochondrial biogenesis and lipid transport (Dixen et al. 2013; Schreiber et al. 2004; Singh et al. 2018), binding of ERR to glycolytic targets has been observed by ChIP-seq in breast cancer cell lines (Cai et al. 2013). In addition, hepatocyte ERRα directly regulates lactate dehydrogenase B expression (Gao et al. 2026), suggesting ERR-dependent regulation of glycolytic genes in other cell types. Considering *Drosophila* has one ERR homolog (compared to three in mammals (Giguere 2008)), it is likely that ERR may have overlapping roles in *Drosophila* adipocytes to regulate energy metabolism.

### ERR directly binds to genes of the pentose phosphate pathway in adult adipocytes

Glycolytic intermediates can be utilized by the pentose phosphate pathway (PPP) for nucleotide production and redox homeostasis (**Figure 3a**) (TeSlaa et al. 2023), and PPP enzymes were enriched in our PANGEA analysis for genes occupied by ERR in adult adipocytes (**Figure 1d**). We therefore examined the specific genes bound by ERR in female adipocytes compared to males. Both males and females showed binding for genes encoding four of eight PPP enzymes (*Pgls*, *Rpe*, *Tkt*, and *Taldo*) (**Figure 3b**). Females showed enhanced binding for *Pgd* compared to males; whereas males showed enhanced binding for *Rpi* and *Prps*. Notably, *Zw* (G6PD in mammals) was not bound by ERR in either females or males. These results suggest that ERR may bind to PPP genes in adult adipocytes to shuttle glycolytic intermediates for nucleotide biosynthesis.

**Figure 3.**
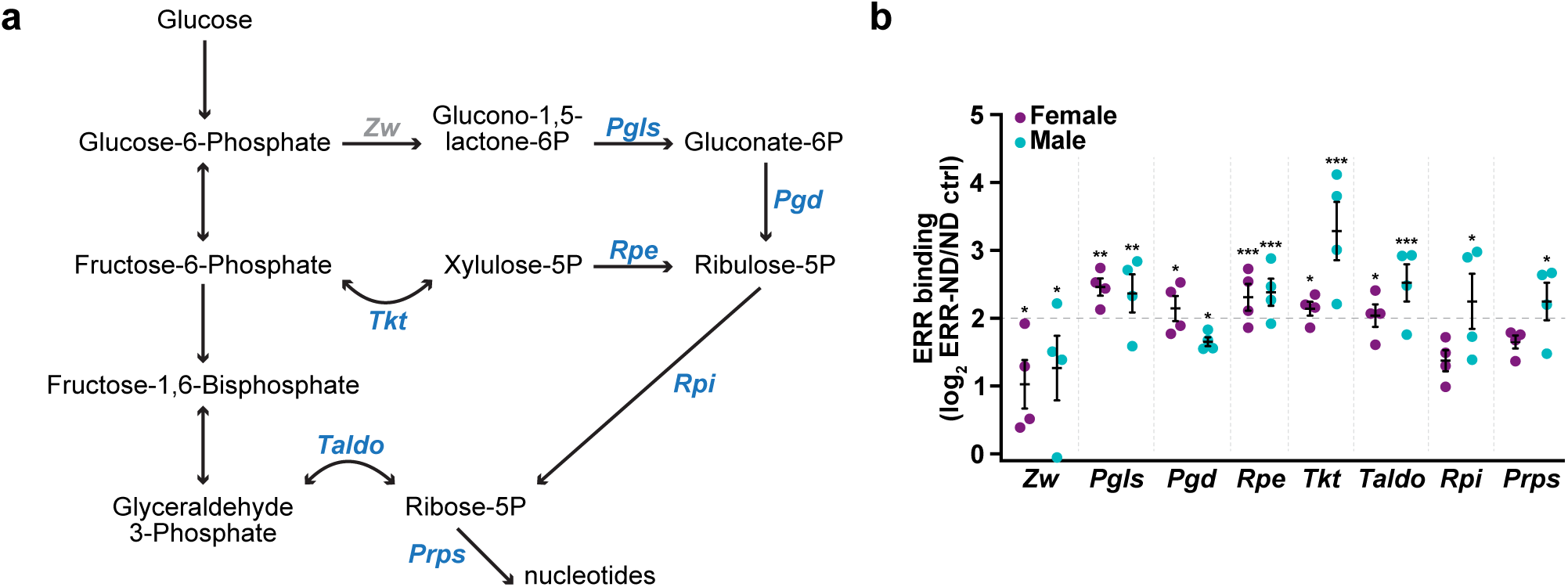
ERR binds to genes encoding pentose phosphate pathway (PPP) enzymes in adult female and male adipocytes. **(a)** Diagram of the PPP shunt showing intermediates and relevant enzymes. Enzymes bound by ERR in adult female or male adipocytes are in bold, blue font. **(b)** Quantification of ERR binding of PPP enzymes in female and male adipocytes. Data from four independent experiments shown as mean ± SEM. Dotted line indicates threshold ratio of ERR-ND/ND ctrl = 2. Values ≥ 2 were considered significant binding as a potential ERR-NanoDam target. \**P* < 1x10^-10^, \*\**P* < 1x10^-15^, \*\*\**P* < 1x10^-20^; Wald test.

Conditional knockout of *ERR* in *Drosophila* adults also decreases PPP transcript expression, with four genes overlapping with those identified by adipocyte ERR-NanoDam in females (*Pgls*, *Pgd*, *Rpe*, *Tkt*) and males (*Pgls*, *Tkt*, *Taldo*, and *Rpi*). Interestingly, *Zw* is downregulated in *ERR* conditional knockouts (Beebe et al. 2020; Fleck et al. 2026), but is not bound by adipocyte ERR-NanoDam, suggesting ERR binds to and transcriptionally regulates PPP genes in other tissues or some PPP genes are not direct ERR targets. While whole body *dERR* mutant larvae showed decreased expression of genes involved in the PPP (Tennessen et al. 2011), fat body RNA-seq analysis of *dERR* mutants showed only significant downregulation for *Rpi* and *Taldo* (Fasteen et al. 2025). These results suggest that all components of the PPP may not be bound by ERR in adipocytes. However, considering these studies were performed from whole body, mixed sex *dERR* mutants, it will be important to determine whether ERR binds to additional PPP components specifically in larval male and female adipocytes. Our results, therefore, suggest that ERR binds to PPP enzymes in adult *Drosophila* adipocytes to possibly regulate glucose catabolism in a tissue-specific manner.

### Gluconeogenesis enzyme genes are bound by ERR in adult adipocytes

Gluconeogenesis is the process of producing glucose from noncarbohydrate substrates (**Figure 4a**) (Shah and Wondisford 2023) and is required for regulating whole body glucose homeostasis. Fly Metabolic Pathway analysis using PANGEA suggests that genes encoding enzymes involved in glucose production were enriched in both female and male adipocytes for ERR occupancy (**Figure 1d**). In both male and female adipocytes, ERR was bound to three of four gluconeogenesis enzyme genes and the gene encoding the trehalose transporter, *Tret1* (**Figure 4b**). ERR did not have significant occupancy at the genomic locus for *Fbp* (the product of which converts fructose-6-phosphate to glucose-6-phosphate) in either male or female adipocytes.

**Figure 4.**
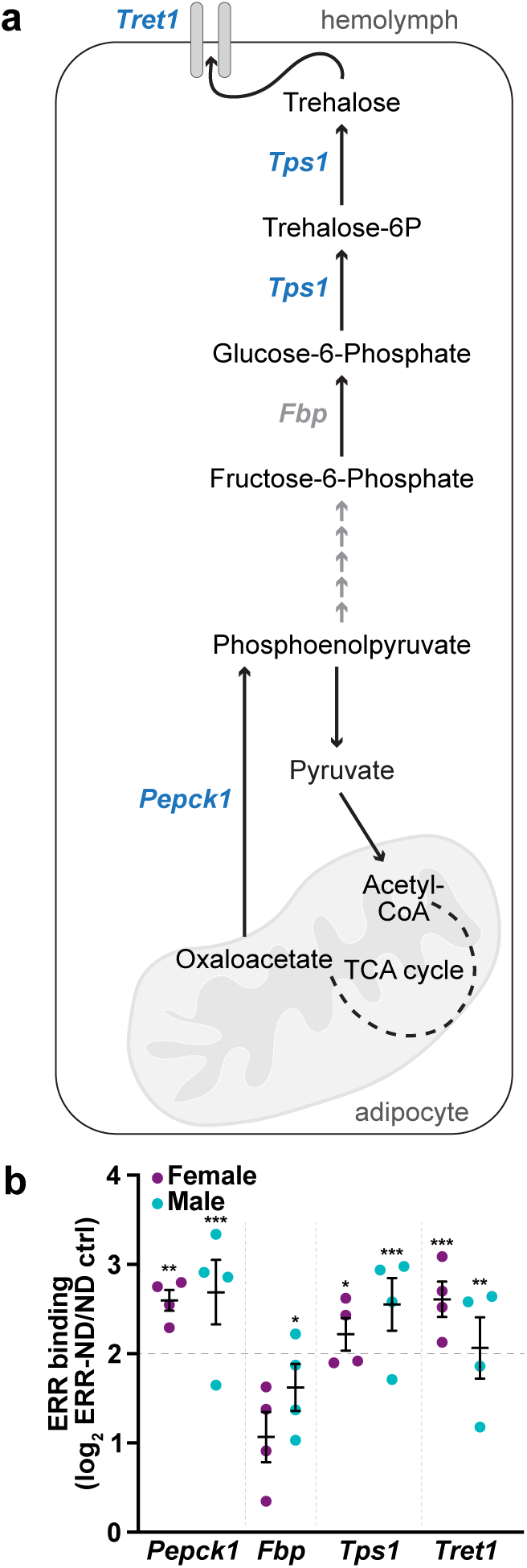
Gluconeogenesis component loci are bound by ERR in adult adipocytes. **(a)** Diagram of the gluconeogenesis pathway and trehalose export through the Tret1 transporter. Genes bound by ERR in male or female adipocytes are bolded in blue. **(b)** Quantification of ERR binding of gluconeogenesis enzymes and the *Tret1* transporter in adult female and male adipocytes. Data from four independent experiments shown as mean ± SEM. Dotted line indicates threshold ratio of ERR-ND/ND ctrl = 2. Values ≥ 2 were considered significant binding as a potential ERR-NanoDam target. \*\**P* < 1x10^-15^, \*\*\**P* < 1x10^-20^; Wald test.

Interestingly, of the targets identified by adipocyte ERR-NanoDam, *Tps1* and *Pepck1* were downregulated only in *ERR* conditional knockout males. This suggests that ERR may be regulating transcription of gluconeogenesis enzyme genes in adipocytes of both sexes, but the inclusion of whole-body transcripts possibly masks differential transcript expression in females. In mammals, glucose homeostasis is controlled by regulation of glucose production in hepatocytes by ERRγ. Hepatocyte-specific loss of *ERRγ* prevents gluconeogenesis induction, whereas *ERRγ* overexpression increases glucose production through induction of *PEPCK* and *G6PC* transcripts (Kim et al. 2012). Conversely, ERRα is a negative regulator of gluconeogenesis, repressing the expression of *PEPCK* transcripts in hepatocyte cell culture (Herzog et al. 2006). While our results suggest that ERR is bound to the loci of these enzymes in adult *Drosophila*, future studies should address whether the activity of ERR is conserved with ERRγ (as an activator) or ERRα (as a repressor) in the regulation of gluconeogenesis.

### ERR binds to β-oxidation enzyme genes in adult adipocytes

β-oxidation occurs in the mitochondria to breakdown fatty acids to acetyl-CoA and requires many intermediate reactions (**Figure 5a**). It was previously shown that *ERR* mutant larval adipocytes have decreased expression of β-oxidation enzymes (Fasteen et al. 2025), therefore we analyzed whether any of these enzymes were bound by ERR in adult male and female adipocytes. Of the enzymes involved, only five of 11 genes (*CG5321*, *CG4335*, *CG3902*, *scully*, and *yip2*) showed binding by ERR in either male or female adult adipocytes (**Figure 5b**). We also noted that four genes (*CG5321*, *CG3902*, *scully*, and *yip2*) bound by ERR in adults were also downregulated in *ERR* mutant larval adipocytes (Fasteen et al. 2025), and two (*GC5321* and *CG3902*) were downregulated in *ERR* conditional knockout males (Beebe et al. 2020). Interestingly, previous RNA-seq analysis from whole body *ERR* mutant larvae or adult *ERR* conditional knockout females failed to identify differentially expressed β-oxidation genes (Fleck et al. 2026; Tennessen et al. 2011), suggesting ERR may generally regulate β-oxidation in adipocytes but some transcript changes cannot be observed from whole body RNA-seq samples. Although it is also possible that ERR does not regulate β-oxidation in females. In addition, association of ERRs with fatty acid oxidation enzymes has been observed in mammals. For example, ERRα is a regulator of medium chain acyl-coenzyme A dehydrogenase (MCAD) (Sladek et al. 1997) and binds to the nuclear receptor response element in differentiated brown adipocytes (Vega and Kelly 1997). Collectively, these results suggest that ERRs can regulate lipid metabolism in adipocytes, but their direct targets may alter depending on the organism and developmental stage.

**Figure 5.**
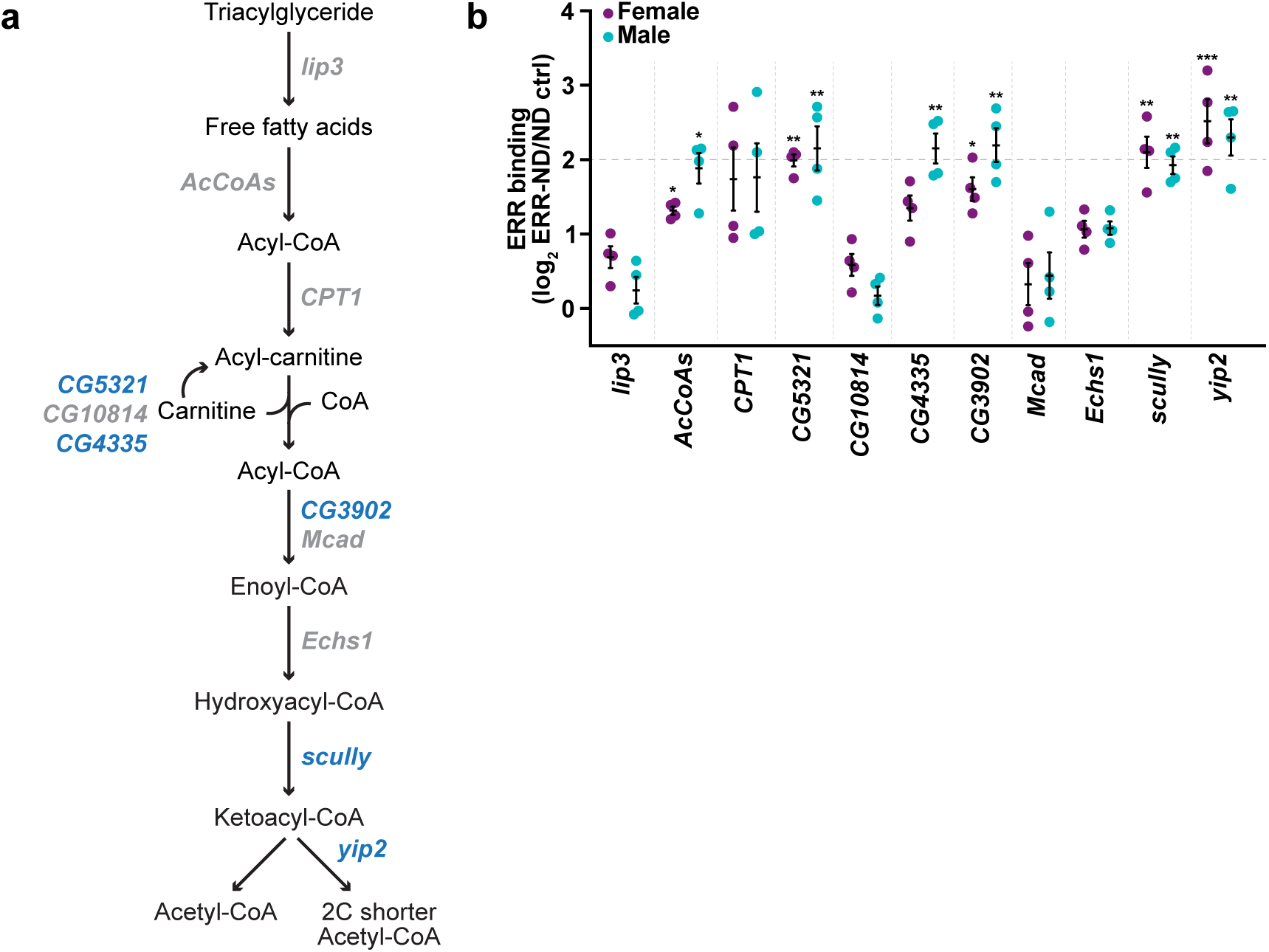
ERR binds to some genes encoding fatty acid oxidation enzymes in adult female and male adipocytes. **(a)** Diagram of fatty acid breakdown and representative enzymes. Genes bound by ERR in adult female or male adipocytes are in bold, blue font. **(b)** Quantification of ERR binding of β-oxidation enzymes in females and male adipocytes. Data from four independent experiments shown as mean ± SEM. Dotted line indicates threshold ratio of ERR-ND/ND ctrl = 2. Values ≥ 2 were considered significant binding as a potential ERR-NanoDam target. \**P* < 1x10^-10^, \*\**P* < 1x10^-15^, \*\*\**P* < 1x10^-20^; Wald test.

Overall, our genomic profiling of ERR in adult male and female adipocytes using NanoDam suggests that components of central metabolic pathways such as glycolysis, the PPP, and fatty acid oxidation are bound by ERR in adult *Drosophila* adipocytes. These results further support that ERRs across species and sexes have conserved features in regulating energy homeostasis. Furthermore, our results highlight that ERR targets are likely sexually dimorphic and may also change depending on the developmental stage in the same organism. Importantly, cell type-specific genomic profiling of ERR in a multi-cellular tissue identified potential direct targets that were not be observed with bulk RNA-seq analysis. Our results show that while ‘omics-style approaches serve as an initial step in identifying the major biology pathways governed by transcription factors, they must be followed by functional analysis and biochemical confirmation. Given the role of ERR in metabolic processes, future studies should also explore the unique ERR targets in different cell types, during different developmental time scales, and under changing physiological conditions.

## DATA AVAILABILITY STATEMENT

The *Drosophila UAS-NanoDam* stocks were obtained from Dr. Andrea Brand and require a Material Transfer Agreement negotiated between University of Cambridge and the researcher’s respective university. Additional *Drosophila* strains are available from the Bloomington *Drosophila* Stock Center (BDSC). The data and analyses in the paper are described in the main text. The raw data and processed data files for adult adipocyte RNA-seq and NanoDam are available through the NCBI GEO accession number GSE341741 and are also provided as supplemental tables.

## ACKNOWLEDGMENTS

*Drosophila* stocks used in this study (except the *UAS-NanoDam* strains) were obtained from the BDSC (NIH P40OD018537). We are thankful to Flybase (flybase.org, NIH 5U24HG013300), an invaluable *Drosophila* research resource. We are grateful to Brian R. Calvi and Deepika Vasudevan for critical reading of the manuscript. This work was supported by the National Institutes of Health R35 GM150517 grant to L.N.W.

